# Profiling microglia from AD donors and non-demented elderly in acute human post-mortem cortical tissue

**DOI:** 10.1101/2020.03.18.995332

**Authors:** Astrid M. Alsema, Qiong Jiang, Laura Kracht, Emma Gerrits, Marissa L. Dubbelaar, Anneke Miedema, Nieske Brouwer, Maya Woodbury, Astrid Wachter, Hualin S. Xi, Thomas Möller, Knut P. Biber, Susanne M. Kooistra, Erik W.G.M Boddeke, Bart J.L. Eggen

## Abstract

Microglia are the tissue-resident macrophages of the central nervous system (CNS). Recent studies based on bulk and single-cell RNA sequencing in mice indicate high relevance of microglia with respect to risk genes and neuro-inflammation in Alzheimer’s disease. Here, we investigated microglia transcriptomes at bulk and single cell level in non-demented elderly and AD donors using acute human post-mortem cortical brain samples. We identified 9 human microglial subpopulations with heterogeneity in gene expression. Notably, gene expression profiles and subcluster composition of microglia did not differ between AD donors and non-demented elderly in bulk RNA sequencing nor in single-cell sequencing.

## Introduction

Alzheimer’s disease (AD), one of the most prevalent age-related neurodegenerative disorders, is characterized by extracellular deposition of β-amyloid protein (Aβ) and intra-neuronal neurofibrillary tangles in the neocortex [1].

Functional changes occurring in microglia cells have been proposed as an important factor in AD pathology [2,3]. AD single nucleotide polymorphism (SNP) heritability was recently found to be highly enriched in microglia enhancers [4]. Multiple genes associated with increased susceptibility for sporadic AD are preferentially expressed in microglia, including *APOE, CR1, CD33, INPP5D, PLCG2, MS4A6A* and *TREM2* [5,6]. In AD mouse models, microglia have been implicated in Aβ seeding, Aβ plaques are surrounded by activated microglia, microglia protrusions physically interact with insoluble Aβ aggregates and microglia around plaques undergo transcriptional changes [7–12]. Sustained depletion of microglia in 5xFAD mice prevents Aβ plaque formation in parenchymal tissue, rather showing Aβ accumulation in the brain vasculature [13]. The functional changes occurring in microglia during AD pathology seem to be diverse [14] and the exact role that microglia play in AD pathology is still unknown.

Many efforts have been made in AD mouse models to identify subpopulations of microglia that are associated with AD pathology. A microglial neurodegenerative subpopulation was discovered by Krasemann and colleagues that was associated with Aβ plaques and triggered by phagocytosis of apoptotic neurons [9]. This transcriptional phenotype was characterized by increased *Spp1, Itgax, Axl, Lilrb4, Clec7a, Ccl2, Csf1*, and *Apoe* and decreased *P2ry12, Tmem119, Olfml3, Csf1r, Rhob, Cx3cr1, Tgfb1, Mef2a, Mafb*, and *Sall1* expression levels [9]. At the same time, a highly similar gene expression change, associated with microglia surrounding Aβ plaques was reported by Keren-Shaul and colleagues [8], termed disease-associated microglia (DAMs). Using single-cell RNA-sequencing these DAMs were subdivided into two sequential stages, a *Trem2*-independent stage, marked by increased expression of *Tyrobp, Apoe*, and *B2m* and decreased expression of homeostatic genes, followed by a *Trem2*-dependent activation stage marked by induction of genes involved in lipid metabolism and phagocytosis (*Trem2, Spp1, Itgax, Axl, Lilrb4, Clec7a, Cts7, Ctsl, Lpl, Cd9, Csf1, Ccl6, Cd68* and more). Sala et al. [15] described a microglia subpopulation that appears in response to Aβ deposition in App^NL-G-F^ mice and shares gene expression changes with DAMs. They identified mutually exclusive subtypes of activated response microglia (ARMs) overlapping with DAMs and, in addition, an independent subtype of interferon response microglia (IRMs).

Studies investigating human microglia subtypes are limited, probably due to the technical and logistical difficulties of isolating pure, viable microglia from acute human brain tissue. Olah and colleagues [16] investigated single human microglia from donors with a large variety of neuropathological backgrounds. They observed 23 clusters of microglia, where 5 out of 23 clusters were enriched for DAM signature genes. However, the neuropathological background of donors was too diverse to associate the observed changes to AD. Mathys and colleagues [17] used single-nucleus sequencing and subclustered ∼2400 microglia of 48 donors. This study was focused on cell-type specific responses to AD development and around 50 microglia per donor was insufficient to fully define microglia diversity in AD.

In this study, we aimed to identify transcriptomic changes in human microglia at the end-stage of AD, by applying both bulk and single-cell sequencing of microglia acutely isolated from acute post-mortem CNS tissue. We isolated and sequenced a pure population of microglia after CD11B+CD45+-based FACS sorting and investigated effects of gender, brain region, tissue dissociation methods and diagnosis.

## Materials and methods

### Human brain specimens

Autopsy brain specimens from the LPS, GFS and lateral temporal lobe were obtained from 25 donors of the Netherlands Brain Bank (NBB, https://www.brainbank.nl/) and 2 donors of the NeuroBiobank of the Institute Born-Bunge (NBB-IBB, Wilrijk, Antwerp, Belgium, ID: BB190113). Mechanical isolation of microglia and bulk microglia sequencing was performed with 22 specimens (11 GFS, 11 LPS) of 15 donors (6 AD, 6 CTR+, 3 CTR). From nine donors both regions were collected and from the other donors either GFS (2016-038, 2016-095, 2017-102) or LPS (2017-098, 2017-124). From three donors (2016-038, 2016-046, 2016-080), LPS or GFS specimens were divided for mechanical as well as enzymatic dissociation. Single-cell sequencing was performed with 16 specimens (15 LPS, 1 temporal) from 15 donors (6 AD, 5 CTR+, 4 CTR), from which 2 were overlapping with donors used in bulk analysis (2018-18, 2018-021). All donors have given informed consent for autopsy and use of their brain tissue for research purposes. The performed procedures, information - and consent forms of the NBB have been approved by the Medical Ethics Committee of the VU Medical Centre. On average, the autopsies were performed within 6h after death. Detailed information about brain specimens used for bulk and single-cell sequencing can be found in Table S1 and S2, respectively.

### Microglia isolation and sorting

The mechanical isolation of microglia was performed as described previously [18,19] with minor modifications. All procedures were performed on ice and all centrifugation steps were performed at 4°C. The tissue was homogenized by mechanical dissociation in Medium A (HBSS (Gibco, 14170-088) containing 15 mM HEPES (Lonza, BE17-737E) and 0.6% glucose (Sigma-Aldrich, G8769) and was filtered through a 300 and 106 µm sieve. Cells were centrifuged at 220g for 10 min and myelin and other lipids were removed through two percoll gradient centrifugation steps. A 100% Percoll solution was prepared consisting of 90% Percoll (GE Healthcare, UK) and 10% 10x HBSS (Gibco, 14180-046), from which the dilutions were prepared. First, cells were resuspended in 24.5% (vol/vol) Percoll in Medium A. A layer of PBS was added and the gradient was centrifuged at 950g for 20 min with reduced acceleration speed and brakes off. After the supernatant was removed, cells were resuspended in 60% (vol/vol) Percoll in 1x HBSS (Gibco, 14170-088) and a layer of 30% (vol/vol) Percoll in 1x HBSS (Gibco,) and PBS, respectively, were added and centrifuged at 800g for 25 min (acc: 4, brake: 0). The cells in between the 30%/60% percoll layer were collected and washed in 1x HBSS (Gibco, 14175-053) and centrifuged at 600g for 10 min.

To examine the effect of the isolation protocol on the microglia expression profile, for three tissue samples (2016-038, 2016-046, 2016-080) mechanical and enzymatic dissociation on two halves of the same tissue was performed. 1.5 gram of tissue was incubated for 80 minutes at 37°C continuously shaking in enzyme solution (1x Trypsin/EDTA (Gibco, A1217701), 0.1 mM HEPES (Lonza, BE17-737E), 0.5 mg/ml DNase I (Roche, 10104159001) in 1x HBSS (Gibco, 14170-088)). Next, it was continued with the Percoll gradient centrifugation steps as described above.

Cells were incubated with anti-human Fc receptor (0,005 µg/ml eBioscience, 14-9161-73) for 10 min in Medium A without phenol red (HBSS (Gibco, 14170-053) containing 15 mM HEPES (Lonza, BE17-737E), 0.6% glucose (Sigma-Aldrich, G8769), 1mM EDTA (Invitrogen, 15575-038)), followed by the incubation with 5 µg/ml DAPI (Biolegend, 422801), eBioscience DRAQ5 (Thermofisher Scientific, 63351), FITC anti-human CD45 (BioLegend, 304006) and PE anti-human CD11B (BioLegend, 301306). Single, viable microglia defined as DAPI^-^, DRAQ5^+^, CD45^+^ and CD11B^+^, were FACS sorted on a Beckman Coulter MoFlo XDP or Astrios.

For bulk microglia RNA sequencing, microglia were sorted into low retention Eppendorf tubes (Sigma, Z666548-250EA) containing 200 µl RNA later (Qiagen, 76104). Following, centrifugation at 4°C and 5000g for 10 minutes, supernatant was carefully removed and microglia were resuspended in 350 µl RLTplus lysis buffer (Qiagen, 1053393) and stored at − 80°C. For barcoded 3’ single-cell sequencing, 15792 single microglia were collected in 384-well PCR plates containing cell lysis buffer (0,2% Triton (Sigma-Aldrich, T9284), 4 U RNAse inhibitor (Takara, 2313A), 10mM dNTPs (Thermo Scientific, #R0193) and 10µM barcoded oligo-dT primer) and were stored for maximally one month at −80°C until further processing. For 10x Genomics Chromium single cell RNA sequencing, approximately 25000 single microglia were sorted from two samples (2018-135, 2ß19-010) into low retention Eppendorf tubes (Sigma, Z666548-250EA) containing 5 µl Medium A and were immediately processed with the Single Cell 3’ Reagent Kit v2 (10x Genomics). FACS data was analyzed with FlowJo (Becton, Dickinson & Company).

### Bulk microglia RNA sequencing library preparation

Total RNA was extracted from the bulk sorted microglia samples using the RNeasy Plus Micro kit (Qiagen, 74034) according to the manufacturer’s protocol. RNA quality and quantity were determined with the Experion RNA HighSens Analysis Kit (Bio-Rad, #7007105). All 25 RNA samples, with RIN values varying between 5.7 and 9.9, were enriched for poly(A)+ messenger RNA using NEXTflex Poly(A) Beads (BIOO Scientific, #NOVA-512980) according to the protocol in the manual. Fourteen µL of this mRNA-enriched poly(A)-tailed RNA was used as the input for the NEXTflex Rapid Directional qRNA-Seq kit (Bioo Scientific, #NOVA-5130-04). Library preparation was performed according to the manufacturers protocol. Quality and concentration of libraries from individual samples were assessed using the High Sensitivity dsDNA kit (Agilent, 067-4626) on a 2100 Bioanalyzer (Agilent) and a Qubit 2.0 Fluorometer (Life Technologies). Subsequently, individual libraries were combined into 2 sequencing pools of 13 samples each with equal molar input. 75bp paired-end sequencing was performed on an Illumina NextSeq 500 system. PhiX was added at 5% to both pools as an internal control before sequencing.

### Single-cell RNA sequencing library preparation

The single-cell RNA library preparation method that was used here is based on the Smart-seq2 protocol by Picelli [20], with the modification of obtaining 3’ instead of full-length RNA/cDNA libraries as in Uniken Venema [21]. After cell lysis and barcoded poly-dT primer annealing (73°C, 3 min), RNA was reversed transcribed (RT) based on the template switching oligo mechanism using 0.1 µM biotinylated barcoded template switching oligo (BC-TSO, 5’-AAGCAGTGGTATCAACGCAGAGTACATrGrG+G-biotin-3’), 25 U SmartScribe reverse transcriptase, first-strand buffer and 2mM DTT (Takara, 639538), 1 U RNAse inhibitor (Takara, 2313A) and 1M betaine (Sigma-Aldrich, B0300-5VL) with the following PCR program: 1) 42 °C 90 min, 2) 11 cycli of 50 °C 2 min, 42 °C 2 min, 3) 70 °C 15 min. To account for amplification bias and to allow multiplexing of cells and samples, the barcoded poly-dT primer contains a cell-specific barcode and a unique molecular identifier (UMI) and a known sequence that is used as a primer binding site during the first amplification step. This same primer binding site is linked to the BC-TSO enabling the use of one primer pair (custom primer) during the first amplification. After the RT reaction, primer-dimers and small fragments were removed by 0.5 U Exonuclease (GE Healthcare, E70073Z) treatment for 1 h at 42 °C. cDNA libraries were amplified with KAPA Hifi HotStart ReadyMix (KAPA Biosystems, KK2602) and custom PCR primer (5’-AAGCAGTGGTATCAACGCAGAGT-3’) with the following PCR program: 1) 98 °C 3 min, 25 cycles of 98 °C 20 s, 67 °C 15 s 72 C 6 min, 3) 72 C 5 min. cDNA libraries of 84 cells were multiplexed and short fragments were eliminated by Agencourt Ampure XP beads (Beckman Coulter, A63880, ratio of 0.8:1 beads to library volume). The quality of multiplexed cDNA libraries was examined with a 2100 Bioanalyzer (Agilent) according to the manufacturer’s protocol. cDNA libraries with an average size of 1.5-2 kb were tagmented and indexed during a second PCR amplification step with the Illumina Nextera XT DNA preparation kit (Illumina, FC-131-1024). Tagmentation was performed according to the manufacturer’s protocol with an input of 500 pg cDNA and amplicon tagment mix for 5 min at 55 °C. The tagmentation reaction was stopped using NT buffer. Next, tagmented cDNA was amplified with Nextera PCR master mix, the Nextera indices (12 pool-specific indices, Illumina, FC-131-2001) and 10 µM P5-TSO hybrid primer (5’-AATGATACGGCGACCACCGAGATCTACACGCCTGTCCGCGGAAGCAGTGGTATCA ACGCAGAGT*A*C-3’) with the following PCR program: 1) 72 °C 3 min, 2) 95 °C 30 s, 3) 10 cycles of 95 °C 10 s, 55 °C 30 s, 72 °C 30 s, 4) 72 °C 5 min). Tagmented cDNA libraries were purified by 0.6:1 ratio of Agencourt Ampure XP beads (Beckman Coulter, A63880) to library volume. The quality and concentration of tagmented cDNA libraries were determined with a 2100 Bioanalyzer (Agilent). cDNA pools with an average size of 300-600 bp were multiplexed using a balanced design with pools from 10 different donors (in total 840 cells) per sequencing run. In other words, cells from each donor were distributed over several sequencing runs to avoid potential batch effects. To eliminate short fragments, the final superpool was gel-purified from 2% agarose gel (Invitrogen, 10135444) with the Zymoclean Gel DNA Recovery kit (Zymo Research, D4007). The concentration was determined using a 2100 Bioanalyzer (Agilent) and Qubit 3.0 (ThermoFisher Scientific) according to the manufacturers protocol. Pools were loaded on an Ilumina NextSeq at a final concentration of 2 pM with7% spike in of PhiX DNA. 0.3 µM BC read 1 primer (5’-GCCTGTCCGCGGAAGCAGTGGTATCAACGCAGAGTAC-3’) was used for the sequencing run. The libraries were sequenced on an Illumina NextSeq 500 system with an average sequencing depth of 17,391 UMIs per cell.

### 10x Genomics Chromium single cell 3’ library construction

The single-cell barcoded libraries were constructed according to the instructions of the Single Cell 3’ Reagent Kits v2 (10x Genomics). Briefly, cells were loaded into a slot of a Chromium chip and GEMs were incubated in a thermal cycler to generate barcoded cDNA. After amplification, the cDNA was fragmented and processed for sequencing by ligating adapters and sample indices. The libraries were sequenced on an Illumina NextSeq 500 system with a sequencing depth up to 20,000 reads per cell.

### Immunohistochemistry

Immunohistochemistry was performed as described previously [11]. Briefly, 16 μm sections of PFA-fixed human brain tissue were vacuum-dried, post-fixated for 10 minutes with 4% PFA, and washed with PBS. Heat-induced antigen retrieval was performed in sodium citrate solution (pH 6.0) for 10 min in a microwave. Endogenous peroxidase was blocked by incubating the slides in 0.3% H2O2 for 30 min. After three washing steps with PBS, primary antibodies against IBA1 (WAKO, 019-19741, 1:1000), Phospho-TAU (ThermoFisher, MN1020, clone AT8, 1:750) and Beta-Amyloid (Dako, M0872, 1:100) were diluted in Bright Diluent (ImmunoLogic, BD09-500) to prevent specific background staining and incubated overnight at 4°C. After three washing steps in PBS, secondary biotinylated horse anti-mouse IgG antibody (0.000125 mg/ml Vector BA-2001) was incubated for 1 hour at room temperature. The tissue sections were washed three times in PBS. The signal was amplified by induction with avidin-biotin complexes (Vectastain Elite ABC-HRP (Vector, PK-6100)) for 30 min at RT and visualized with 3,3′-diaminobenzidine (Sigma, D-5637). Additionally, after the phospho-TAU staining we performed a crystal violet counterstaining. The slides were dehydrated with an ethanol series (50%, 70%, 80%, 90%, 2x 96% and 3x 100% ethanol) and air-dried for 30 min prior to mounting a coverslip with DePeX (Serva, 18243). Imaging was performed with a Hamamatsu Nanozoomer at a 40x magnification.

### Preprocessing of RNA-sequencing data

For bulk samples, the first preprocessing step was stripping the NEXTflex barcode (9 base pairs) from the sequence. The barcode was saved in a separate file for further use. Next, microglia bulk sequencing reads were aligned with HISAT2 (version 2.1.0) to the GRCh38.92 reference genome with Ensembl annotation [22]. Further processing was done with samtools (version 1.9) and Picard Tools (version 1.140). This included sorting, assigning reads to a read group, verification of mate pair information and marking duplicates. After these processing steps, reads were quantified using featureCounts (Subread version 1.6.2) [23]. Reads from bulk samples were deduplicated using a bash script by BIOO Scientific (v2, date 11/1/14), using the NEXTflex barcode that was saved in the additional file. This allows proper elimination of PCR duplicates taking advantage of stochastic labeling of UMIs.

Reads from bc-Smart-seq2 single cells were demultiplexed based on cell barcodes using UMI-tools (version 0.5.3) [24]. We created a whitelist of cell barcodes by applying the *whitelist* function of UMI-tools. Next, the whitelist is compared to the true cell barcodes, and these barcodes are saved as a cell barcode list that is used for downstream processing. The UMI-tools *extract* function was used to extract the cell barcode and the UMI from each read and add this information to the read name of the sequence. Reads were single-end aligned with HISAT2 (version 2.1.0) to the GRCh38.91 reference genome with Ensembl annotation using the default parameters, followed by sorting and indexing of the BAM files.

Primary counts were quantified with the function featureCounts (Subread version 1.6.0) using the flag –primary and -R BAM to save the BAM file. PCR duplicates were removed and unique molecules were counted per gene and per cell using the function UMI-tools *count* [24]. Seventeen pools with less than 10% of reads assigned to genes were excluded.

Reads from 10x Genomics Chromium single cells were demultiplexed and aligned to the GRCh38 genome with Ensembl transcriptome annotation with Cell Ranger using default parameters. The demultiplexed fastq files were used as input for the 10x Genomics pipeline Cell ranger. Barcode filtering was performed with the R package DropletUtils with a threshold of > 100 UMIs per barcode [25].

### Downstream analysis

For bulk samples, DAFS filtering was used to remove lowly expressed genes [26] and principal component analysis was computed on rlog transformed counts using the DESeq2 R package (v1.24.0). We applied upperquartile normalization to adjust for library size and used edgeR (version 3.26.8) for differential gene expression analysis. Average library size after filtering was 3.9M counts (± standard deviation 1.9M). To model gene expression levels we used a negative binomial generalized log-linear model as implemented in edgeR, adjusting for gender, analyzing different brain regions separate. Differences between groups were tested with likelihood ratio tests. Thresholds were set at FDR > 0.05 and abs(logFC) > 1 to define DEGs. For bc-Smart-seq2 single cells, approximately 25% of the cells were filtered out during preprocessing. Downstream analysis started with 16 samples and 12,861 single cells with detectable cell barcodes from 15 different donors. One donor (iB6399-BA7) contributed tissue from both the temporal cortex and the superior parietal cortex. Cell library sizes before filtering were quite different across donors; this is most likely due to differences in tissue quality. To remove empty cells while respecting variation across donors, we set a threshold for each donor individually, removing cells with library sizes exceeding median of log(total counts) ± 3 median absolute deviation (mad) [27]. In addition, cells with more than 3000 unique genes were considered doublets (genes per cell median 520; mad ±276) and cells with >10% mitochondrial reads were excluded. We excluded ∼10% of the cells (differs per donor) based on these criteria. In total, this left 11,419 cells for analysis. Genes not detected in at least 3 cells were removed. After filtering, we observed a median of 13,394 UMIs and median 520 genes per cell for bc-Smart-seq2 data.

For 10X data, similarly, low quality cells with >10% mitochondrial gene (MT) content were removed in donor 2018-135. Donor 2019-010 had very high cell quality, so a >5% MT threshold was applied. Duplicate cells were filtered by setting an upper UMI threshold that was based on the multiplet rate as mentioned in the 10x Genomics user guide. We analyzed 3,077 single cells for MCI donor 2019-010 and 2,881 single cells for AD donor 2018-135.

Raw counts were normalized by total expression per cell, scaled by 10,000, and log-transformed with the CRAN package Seurat (v3.0.0). We used the mean variability method to select highly variable genes (HVGs). Briefly, this method identifies variable genes while controlling for the strong relation between gene variability and gene average expression. We allowed lowly expressed genes in the highly variable gene list, since some disease associated genes (e.g. TREM2, TYROBP, CTSD) are biologically relevant but also lowly expressed. These extra ∼632 lowly expressed HVGs did not change clustering results and were included in the final clustering analysis. Unwanted sources of variation, such as number of detected genes, ribosomal content, and mitochondrial content were regressed out. We used the first 20 principal components for clustering analysis applying PCA-Louvain maximum modularity clustering as implemented by Seurat. Cluster resolution was set at 0.5, since it formed the most stable plateau in cluster number when considering resolutions from 0.1 to 2.0.

10X single cells were analyzed identically. Each donor was analyzed individually and cluster resolution was adjusted due to higher cell numbers. For donor 2019-135 cluster resolution 0.6 and for donor 2019-010 cluster resolution 0.8 was set. Marker genes between clusters were identified using MAST implemented in Seurat’s *FindAllMarkers* function with default thresholds (logfc.threshold = 0.25, min.pct = 0.1). Gene ontology (GO) analysis on upregulated marker genes was calculated using the ClusterProfiler (v3.12.0) [27] with p-value cut-off of 0.01 and q-value cut-off of 0.05. For FDR control we used the R package q-value (version 3.1.1).

### Gene set analysis

Briefly, raw expression values were normalized to counts per million and log(cpm + 1) transformed. The gene set score was an average over all genes in the set per cell. We compared if, on average, cells in any one cluster showed a higher mean expression of the gene set than all other cells by using multiple linear regression with gene set score as dependent variable and cell library size, donor, and cluster as independent variables. Afterwards, p-values were adjusted with a Bonferroni correction. Visualizations were made with the R package ‘ggplot2’.

## Results

### Isolation of pure microglia from acute post-mortem brain tissue

To investigate transcriptomic changes in microglia during AD, bulk and single-cell sequencing were performed. Post-mortem tissue samples of the superior parietal lobe (LPS), superior frontal gyrus (GFS) and lateral temporal lobe were obtained from 27 donors. The samples were classified into 3 experimental groups based on a clinical diagnostic report provided by the Netherlands Brain Bank/NeuroBiobank Born-Bunge and immunohistochemical analysis of Aβ and hyperphosphorylated tau (PHF-tau): CTR (no dementia, absence of Aβ plaques and PHF-tau), CTR with plaques (CTR+) (no dementia, presence of Aβ plaques and/or PHF-tau) and AD (dementia, AD diagnosis, presence of Aβ plaques and/or PHF-tau). One donor diagnosed with mild cognitive impairment (MCI) and presence of Aβ and PHF-tau plaques was included as well. Representative images of Aβ and PHF-tau immunohistochemical stainings are available in Figure S1. The goal of the stratification between CTR and CTR+ was to ensure that the CTR group is free of undiagnosed AD donors. Microglia were isolated from enzymatically or mechanically dissociated tissue and purified by fluorescence-activated cell sorting (FACS), based on single, viable CD11B and CD45 positive cells. Twenty-five samples (12 LPS and 13 GFS; 22 mechanically and 3 enzymatically dissociated) from 14 donors were sorted and sequenced as bulk samples and 16 samples (15 LPS, 1 temporal lobe) from 15 donors were single-cell sorted and sequenced (bc-Smart-seq2 and 10x Genomics) (Fig 1A). Two donors were included in both the single-cell and bulk cohort. The employed strategy resulted in a pure microglia population, as confirmed by expression of known microglia genes, that were based on previously published adult human microglia [19] and human CNS nuclei [28] gene expression profiles (Fig 1B).

**Figure 1.**
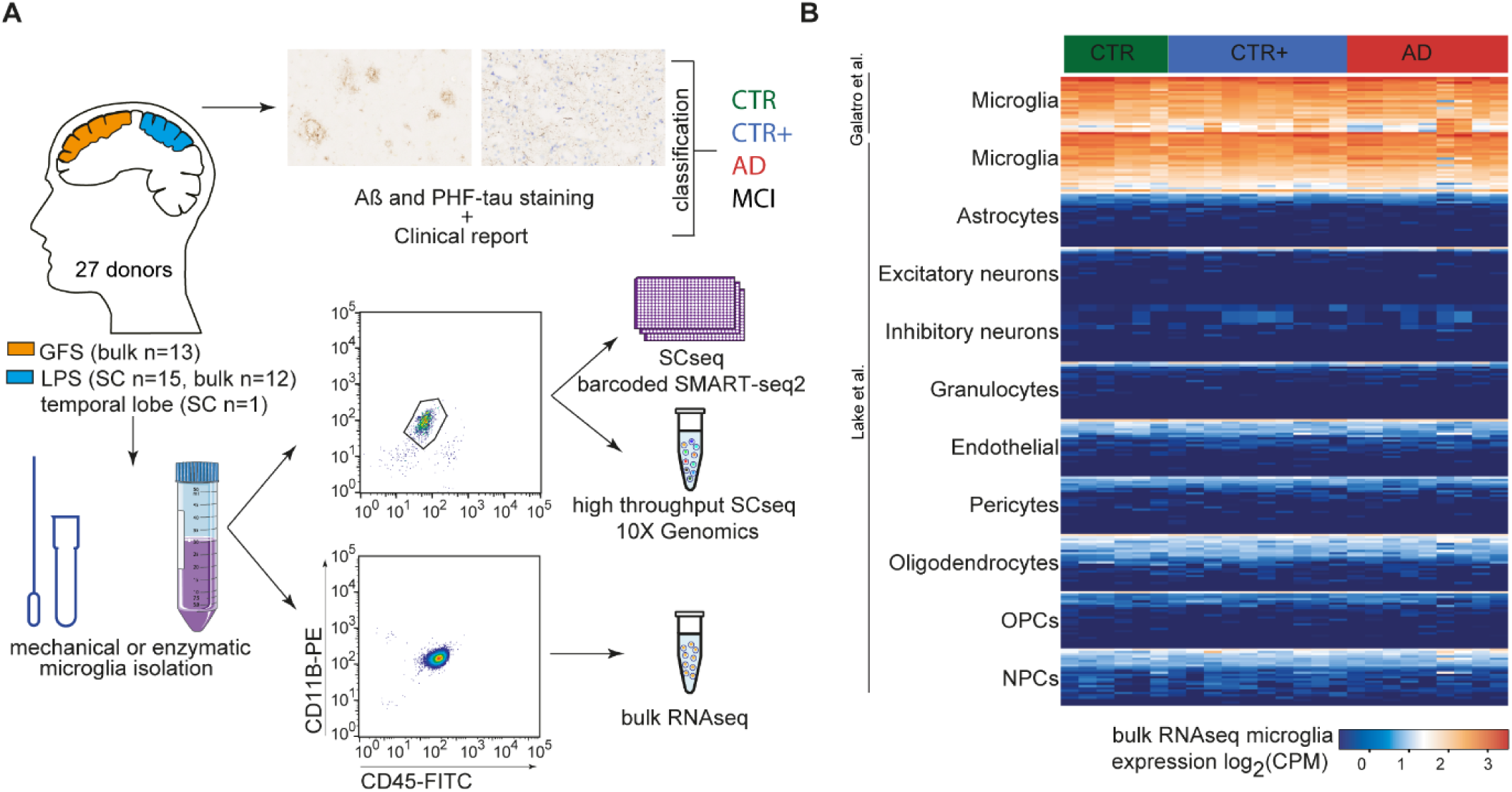
Microglia gene expression profiling of four groups of donors from acute post-mortem human brain tissue. (A) Tissue samples from GFS and LPS were classified into four experimental groups (CTR, CTR+, AD, MCI) based on Aß and Tau immunohistological stainings and the clinical report of the donor. Microglia were mechanically or enzymatically isolated and collected by CD11B+CD45+-based FACS sorting. Sorted microglia were used for bulk or single-cell sequencing (barcoded Smartseq2 and 10x Genomics). Scale bar= 50µm. (B) Expression of top 25 previously published celltype-specific marker genes in bulk-sorted microglia [17, 18]. Abbreviations Aß: amyloid beta, HPF-tau: hyperphosphorylated tau, AD: clinically diagnosed Alzheimer’s Disease with Aβ plaques and/or hyperphosphorylated tau, CTR: Control, CTR+: Control with Aβ plaques and/or hyperphosphorylated tau, GFS: superior frontal gyrus, LPS: superior parietal lobe, MCI: mild cognitive impairment, NPC: Neural Progenitor Cells, OPC: Oligodendrocyte Precursor Cells, SC: single cell.

### No transcriptomic differences between CTR, CTR with plaques (CTR+) and AD donors in acute post-mortem tissue by bulk RNA-sequencing

To investigate general transcriptional characteristics of microglia in AD, we analyzed bulk sorted and sequenced microglia from CTR, CTR+ and AD samples. Principal component analysis (PCA) showed no segregation between donor groups (Fig 2A). The second principle component showed segregation of microglia samples on gender, but not on brain region or age (Fig 2A).

**Figure 2.**
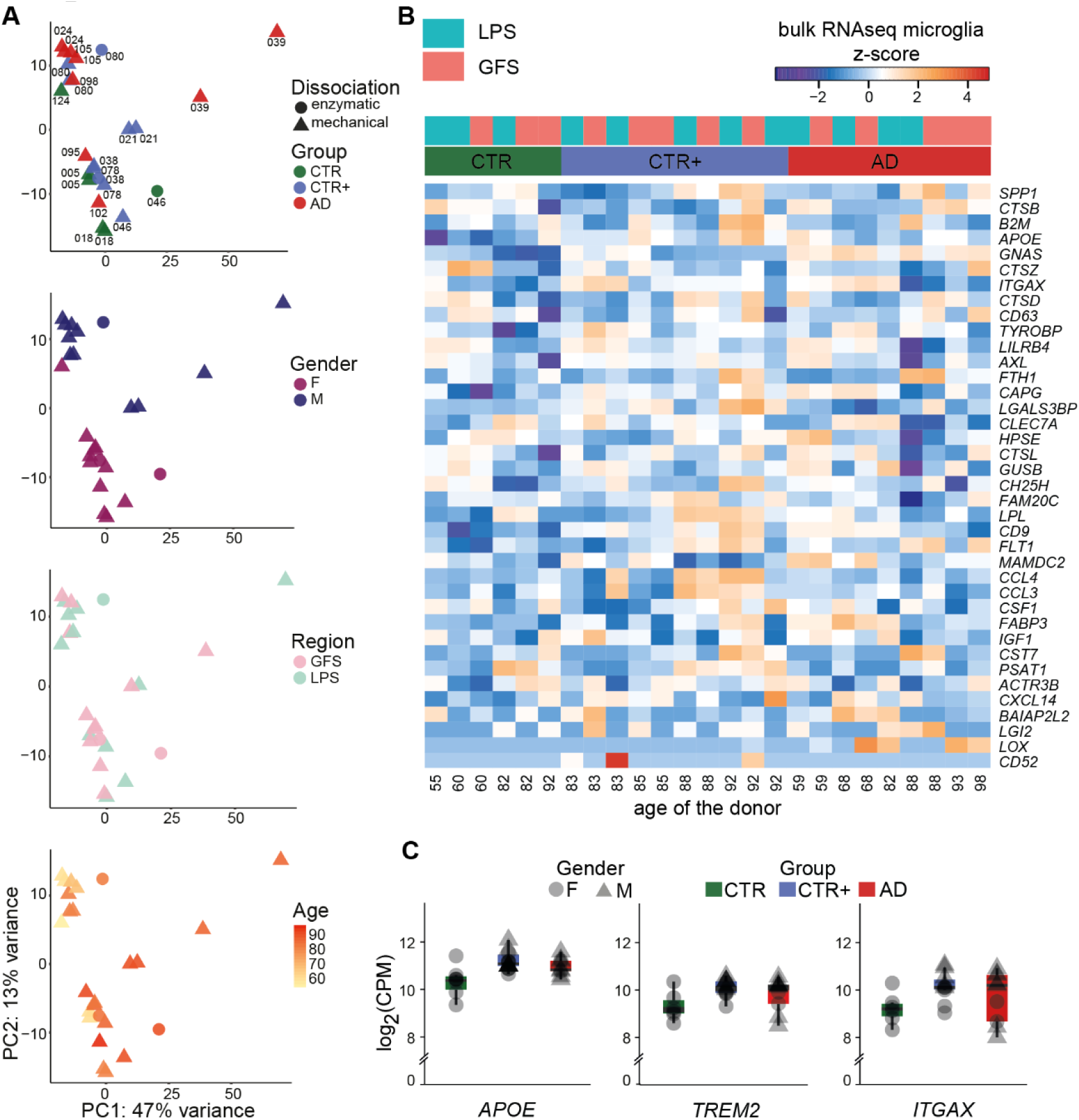
Transcriptomic analysis of microglia populations isolated from CTR, CTR+ and AD donors. (A) PCA of RNA-sequencing data from post-mortem isolated microglia illustrating the effect of tissue dissociation method, donor (each sample is indicated with the donor label), donor groups, gender, brain region, and age. (B) Expression of disease-associated microglia genes in bulk microglia samples from three donor groups (z scores). Donors are ordered based on age within their group. (C) Selected examples of disease-associated microglia gene expression (log2(CPM)) in bulk microglia samples. Abbreviations: AD: Alzheimer’s Disease, CPM: counts per million, CTR: Control, CTR+: Control with Aβ plaques and/or hyperphosphorylated tau, F: female, M: male, GFS: superior frontal gyrus, LPS: superior parietal lobe.

To further examine the effect of gender, male and female GFS microglia samples were compared. Here, 33 genes were differentially expressed with an absolute log fold change > 1, FDR-value < 0.05. One third of these differentially expressed genes (DEGs) were X or Y-linked genes (Table S3).

Differences between microglia from frontal and parietal brain regions were assessed. For 9 donors, microglia were isolated from both the LPS and GFS region, and a within-subject comparison was applied. Only the expression of *JAML* and *TREM1* was significantly higher in LPS compared to GFS (Table S4). This indicates that the gene expression profiles of microglia isolated from frontal and parietal brain regions were very similar.

The effect of the dissociation method on gene expression was determined by comparing the bulk RNA-sequencing profiles of GFS microglia from enzymatically and mechanically dissociated tissue of donors 2016-038 and 2016-080 (4 samples). 10 genes were significantly higher expressed in microglia isolated using enzymatic dissociation: *CXCL8, SOCS1, NR4A1, ID1, FOSB, ATF3, NR4A2, EGR2, OSM* and *HSPA7* (Table S5). As the enzymatic dissociation procedure induced expression of these 10 genes in the bulk samples of these 2 donors, only mechanical dissociation was used for single-cell sequencing.

Bulk samples from AD and CTR donors were compared while correcting for the effect of gender on gene expression. LPS- (n=11) and GFS-derived (n=11) microglia were analyzed separately. In the analysis of microglia from both LPS and GFS regions no differentially expressed genes from AD donors compared to CTR donors were found.

Lastly, expression levels of the top 50 genes upregulated in DAMs [8] were investigated in bulk microglia. These mouse top DAM genes were not differentially expressed between AD-derived compared to control microglia in LPS and GFS regions (Fig 2B). For several known DAM genes, such as *APOE, TREM2, ITGAX* a small increase in expression in AD donors compared to CTR can be recognized, albeit it is not statistically significantly (Fig 2C). To summarize, in bulk transcriptomes of microglia from AD and CTR donors, no significant gene expression differences were detected.

### Single-cell gene expression profiling identifies 9 subsets of microglia but no AD-associated subpopulation

As AD-associated changes possibly only occur in a small subpopulation of plaque-associated microglia and thus might be masked by bulk RNA-sequencing, we next employed single-cell sequencing to investigate microglia heterogeneity. A median of 753 microglia per donor from 4 CTR, 5 CTR+, a donor with mild cognitive impairment (MCI) and 5 AD donors were analyzed. The read depth per cell for each donor was comparable, however the median number of detected genes for each donor varied (Fig S2A,B). Clustering analysis identified 9 microglial subsets, indicating heterogeneity in microglia transcriptomes (Fig 3A). Sequencing depth, uniquely detected genes and detected mitochondrial genes across clusters are visualized in Fig S2. The expression of cell type-specific markers across all clusters showed that all analyzed cells were microglia, and that this population was free from neuronal, erythrocyte, and other glial cell contaminations (Fig 3B). Figure 3C depicts the percentage of cells from a certain donor that contribute to each cluster. Except for cluster 3, all clusters exhibited roughly equal donor contributions. Cluster 3 has a high contribution from AD donor 2018-077 and cells from this donor had a relatively low fraction of mitochondrial genes (Fig S2C) compared to other cells. Gene expression in cluster 3 will strongly depend on this particular donor. Therefore, cluster 3 (9.2% of the cells) was excluded from further analysis.

**Figure 3.**
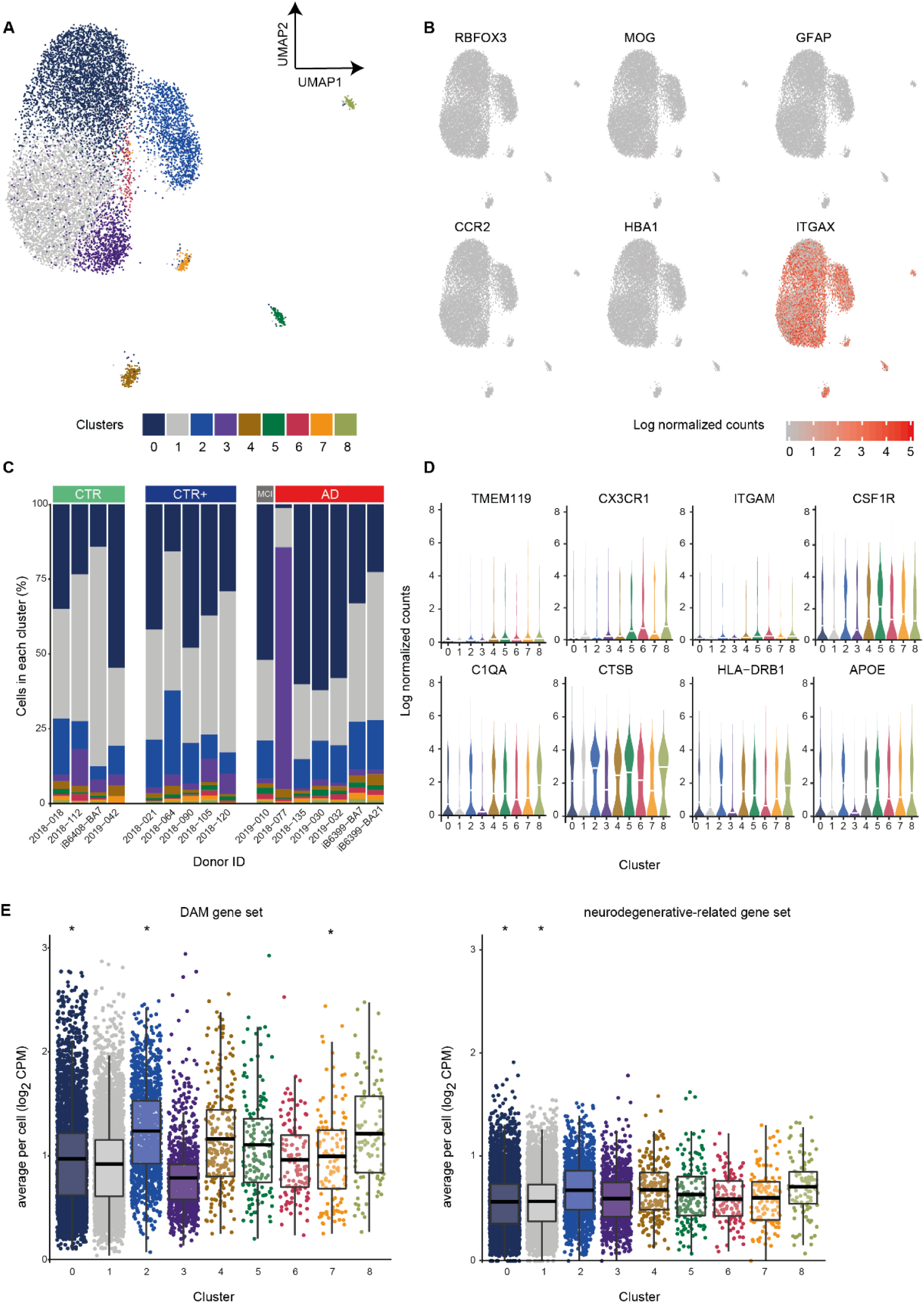
Single-cell expression profiling of microglia identified 9 subsets of microglia but no AD-associated cluster using bc-Smart-seq2. (A) UMAP visualization and unsupervised clustering of microglia derived from AD (n=5), MCI (n=1), CTR+ (n=5) and CTR (n=4) donors. Each dot represents a cell, colors represent cluster identity. (B) Expression of cell type-specific markers for neurons (*RBFOX3*), oligodendrocytes (*MOG*), astrocytes (*GFAP*), infiltrating immune cells as monocytes (*CCR2*), erythrocytes (*HBA1*) and microglia (*ITGAX*). (C) Cluster contribution normalized to donor. Bar height indicates the percentage of cells from a donor that contributed to the clusters. (D) Violin plots show the probability density distribution of normalized gene expression across clusters for selected genes. (E) The average mouse DAM gene set expression in each cell across clusters (left). The average human myeloid neurodegenerative-related gene set expression in each cell (right). Abbreviations: AD: Alzheimer’s Disease, CPM: counts per million, CTR: Control, CTR+: Control with Aβ plaques and/or hyperphosphorylated tau, DAM: disease-associated microglia, UMAP: uniform manifold approximation and projection

Differential gene expression analysis identified marker genes that were significantly enriched in each cluster compared to all other clusters. However, marker genes significantly enriched in a cluster (Table S6) and the biological annotation of those genes (Fig S3) did not point towards clear biological functions of identified clusters. Expression of homeostatic genes such as *CSF1R, ITGAM, TMEM119* and *CX3CR1* was quite variable between cells in a cluster (Fig 3D). These genes were not identified as marker genes in any of the clusters (Table S6). Many microglial genes previously associated with antigen presentation (*HLA* genes), the complement system (*C1QA*), neurodegenerative diseases (*TREM2, TYROBP, B2M*), lysosomal proteases (*CTSB*) and phagocytosis (*CD14, AXL)* were generally expressed at higher levels than homeostatic genes and were found to be marker genes of cluster 2 (Fig 3D).

Unexpectedly, cluster 2 was not enriched by AD-derived microglia and overall no AD-associated microglia cluster was detected. *APOE* expression is associated with AD and was elevated but not significantly enriched in cluster 2 according to differential expression analysis (Fig3D, Table S6). The majority of AD-derived microglia contributed to cluster 0. Cluster 0 was comprised of 58% AD and 42% CTR/CTR+ microglia (Fig 3C).

The small clusters 4-8 contain relatively few cells (0.8-1.8%), they showed increased expression of genes involved in protein degradation such as *PSMD11, PSMB2, PSMB3, EDEM3, FOXN2, UBQLN1* and *DERL2*. Possibly, the small microglia clusters 4-8 reflect cell states with increased stress responses.

In 5XFAD mice, disease-associated microglia (DAMs) were characterized by expression changes in set of genes [8]. To investigate if the increased gene expression in DAMs was also present in one of the identified clusters, the average DAM gene set expression per microglia was calculated. The average DAM gene expression level per cluster was compared with all other clusters. Clusters 0 and 7 showed significantly lower average DAM gene expression levels and microglia in cluster 2 showed significantly higher average DAM gene expression levels (S Table 7), p < 0.001. Although statistically significant, these changes are likely too small to be biologically relevant. These observations were confirmed in a different gene set from a meta-analysis of multiple amyloid mouse models [14]. Again, microglia in cluster 0 showed a significant decrease in neurodegenerative-related gene expression levels. To summarize, only the decrease in gene expression in cluster 0 was consistent and no cluster was associated with increased neurodegeneration-related gene expression (Fig 3E).

Taken together, single-cell analysis of approximately 11,419 microglia which were CD11B+CD45+-based FACS sorted and profiled with bc-Smart-seq2, identified microglial heterogeneity, but no AD-specific microglia subpopulation.

### Microglia diversity but no DAM-like cluster in an individual MCI and AD donor with high microglial cell numbers

The proportion of DAM-like microglia in human AD brain might be relatively low and could potentially be missed by bc-Smart-seq2 expression profiling. Therefore, two donors with ∼3000 microglia each were investigated using the 10X Genomics platform. First, 2881 cells were analyzed from post-mortem LPS tissue of a female AD donor, 81 years old, with high Aβ burden and modest but visible PHF-tau protein (2018-135) (Fig S1). Second, 3077 single cells from an MCI donor (2019-010), 77 years old, with moderate Aβ pathology and high levels of PHF-tau protein in the LPS were analyzed (Fig S1). We applied an identical clustering procedure as for bc-Smart-seq2 libraries, now analyzing each donor individually, only adjusting the clustering resolution for higher cell numbers. Per donor, three clusters were identified (Fig 4A, B). As clustering is performed within one sample, the identified clusters are not confounded by post-mortem delay, gender, tissue quality or other donor-associated factors. Despite testing multiple cluster resolutions, a DAM-like cluster could not be detected.

**Figure 4.**
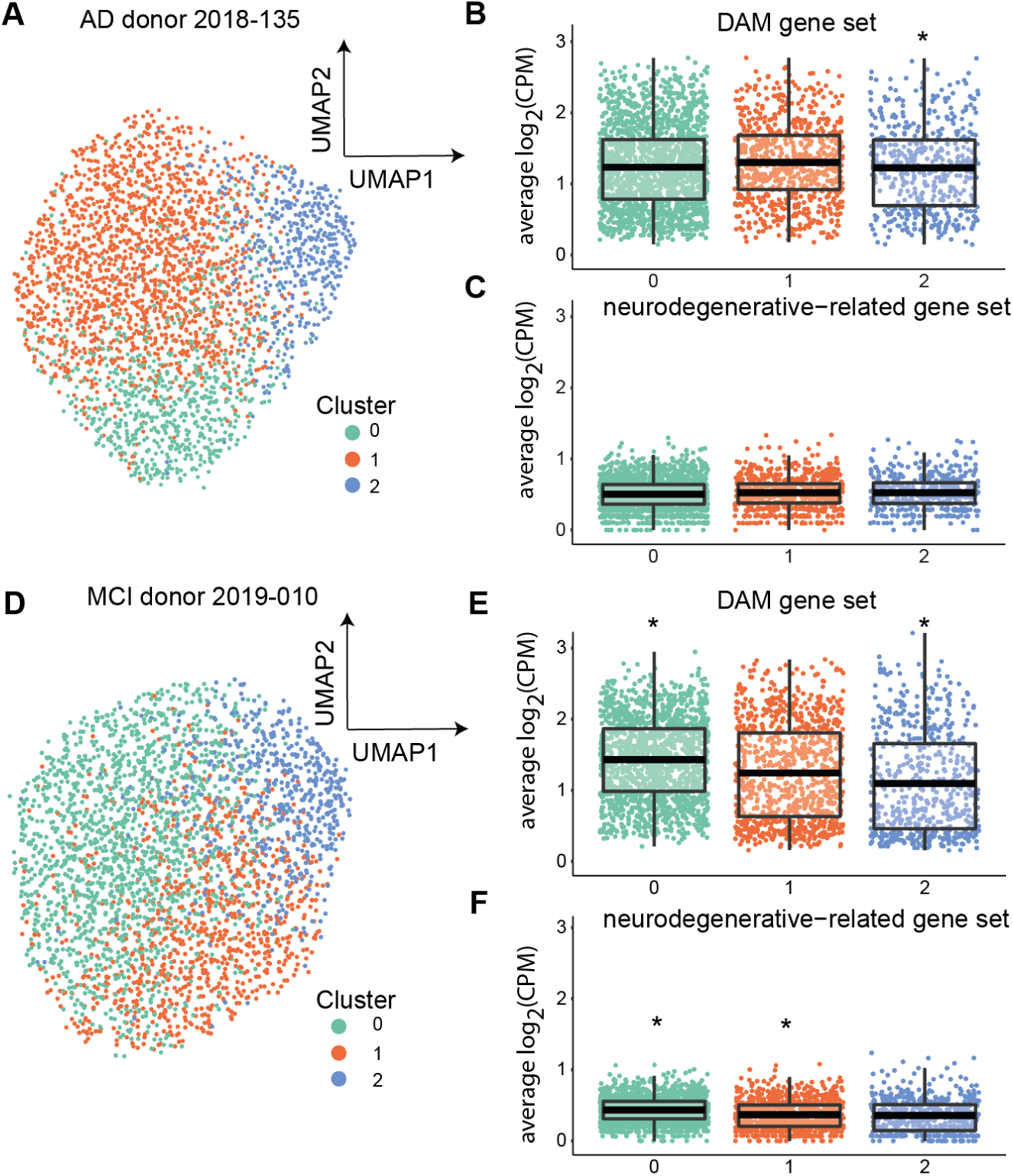
Single-cell expression profiling of microglia from individual donors with high cell numbers using 10X Genomics. (A,B) Unsupervised clustering identified 3 subsets of microglia in AD donor 2018-135 (A) and in MCI donor 2019-010 (D). The average DAM gene set expression per cell and the myeloid neurodegenerative-related gene set per cell across clusters for AD donor 2018-135 (B, C) and for MCI donor 2019-010 (C, F). Abbreviations: AD: Alzheimer’s Disease, CPM: counts per million, CTR: Control, CTR+: Control with Aβ plaques and/or hyperphosphorylated tau, DAM: disease-associated microglia, UMAP: uniform manifold approximation and projection

As described for figure 3, the average gene expression levels were calculated for each cluster and compared to all other clusters. The gene expression averages per cluster were calculated for the mouse DAM gene set [8] and for the neurodegenerative-related gene set [14] (Fig 4B, C, E, F). A significant increase of the DAM gene set and the neurodegenerative gene set was observed in microglia in cluster 0 in the MCI donor (Fig 4 E,F), but not in the AD donor (Fig 4 B,C). In short, no cluster with DAM-like microglia was detected in either donor, although a high number of microglia was analyzed (∼3000 cells/donor).

## Discussion

In this study we aimed to identify transcriptomic changes in human microglia at the end-stage of AD by applying both bulk and single-cell RNA sequencing of microglia isolated from acute post-mortem CNS tissue. In acute parietal and frontal cortex, we analyzed microglia as bulk samples allowing the most sensitive detection of small gene expression changes. Here, transcriptomic differences between AD and CTR were not detected and disease-associated genes identified in AD mouse models were not enriched in AD microglia. Single cell sequencing analysis was applied to detect microglial subtype populations, possibly consisting of low cell numbers, that might be unnoticed in the average transcriptome obtained by bulk analysis. Also in single microglia transcriptomes, the relative contribution to microglia clusters did not differ between AD and control donors, when using the bc-Smart-seq2 protocol on CD11B and CD45 sorted cells.

Expression of the average neurodegenerative-related gene sets were not significantly increased in any cluster. The neurodegenerative disease-associated microglial subtype originally described by Krasemann et al. [9] and Keren-Shaul et al. [8] represented a relatively small fraction of the total microglia population. To rule out that the lack of a DAM-like cluster in our data was due to the analysis of too low cell numbers, a higher number of microglia from two donors were single-cell sequenced using the 10X Genomics platform. Clustering was performed per individual donor to avoid donor variation which might mask a potential DAM cluster. Concordant to the bc-Smart-seq2 data, the top DAM genes were expressed to a similar extent in all clusters. In conclusion, microglial transcriptomes between AD and CTR donors did not differ and a DAM-like microglial subtype was absent in AD donors.

When comparing the clustering results of the 10X Genomics and bc-Smart-seq2 single-cell RNA sequencing techniques, differences were observed. Clusters 0-2 in the bc-Smart-seq2 data contain the vast majority of microglia and are most similar to the three clusters identified per donor in the 10X Genomics data set. The smaller clusters in the bc-Smart-seq2 data (4-8) were not identified in the microglia profiled by 10X Genomics and are possibly associated with the plate-based protocol as we observed similar small clusters in a different bc-Smart-seq2 dataset as well (unpublished results).

There are multiple possible explanations for the absence of AD-associated changes in acute bulk and single cell microglia. The lack of detectable AD-associated changes could be due to limited sample size and donor variation. Another explanation for the lack of AD-related transcripts in bulk and single cell microglia could be that the relevant microglia associated with AD plaques are more vulnerable to the isolation procedure. This would imply that, using conventional isolation and sorting of microglia would enrich a population of cells that are not related to AD pathology. Streit and colleagues first introduced the concept of dystrophic microglia that occur around neuronal structures positive for hyperphosphorylated tau protein [29,30] and that can occur around Aβ plaques as well [31]. Possibly, dystrophic microglia and microglia embedded inside the Aβ plaque are more vulnerable and therefore preferentially lost during FACS gating of live, single cells.

When comparing AD mouse models to human AD, the distinction between parenchymal and plaque-associated microglia might be more pronounced for amyloid mouse models than for human end-stage AD samples. In transgenic amyloid mouse models, especially 5xFAD mice, Aβ is over-expressed in a non-physiological manner, resulting in very fast Aβ plaque formation. Transgenic mouse models lack regional brain atrophy and show less widespread neurodegeneration than human AD cases [32]. Possibly, the parenchymal microglia and the rest of the CNS are less affected in amyloid mouse models. Furthermore, quantifying the cortical dense-core plaque burden in 5XFAD and APPswePS1ΔE9 mouse models showed that in mice, the plaque burden eventually reaches 10 times higher levels than in human AD brain tissue [33]. Additionally, inter-individual variation will influence human microglia transcriptomes more than mouse microglia transcriptomes. Together these observations might lead to a more pronounced change between parenchymal and plaque-associated microglia in amyloid mouse models than in human AD samples. For this reason, single human microglia studies will most likely require much larger cell numbers to capture sufficient plaque-associated microglia than studies using AD mouse models.

DAMs were not only associated with neurodegenerative diseases, but also with natural ageing [8,9]. This was confirmed by Sala and colleagues investigating an App^NL-G-F^-associated microglia subpopulation, ARMs, which overlap with DAMs. ARMs already comprised a few percent of microglia in the brains of wild-type mice at young age and evolve naturally with ageing [15]. Furthermore, a consensus gene expression network module co-occurring both during ageing and neurodegeneration was detected. The described module included DAM signature genes such as *Csf1, Spp1, Apoe, Axl, B2m, Ctsz, Cd9, Cstb*, and *Cst7*. Biological annotation of module-specific genes included phagosome and lysosomal pathways [34]; functions previously associated with DAMs [8]. Taken together, this suggests the presence of DAM-like microglia could be expected, albeit at low levels, in aged controls as well as AD donor-derived microglia.

It is still an unresolved question whether a subpopulation resembling DAMs exist among human microglia. Three other studies previously addressed this question. Olah et al. [16] observed 23 clusters of human microglia, where 5 out of 23 clusters were enriched for DAM signature genes. Three of the 15 donors suffered from AD pathology, making it difficult to connect their microglia subpopulations with AD-induced gene expression changes. Srinivasan et al. [35] investigated frozen myeloid cells from AD brain tissue and observed that from the 100 DAM genes, only expression of *APOE* did indeed change in myeloid cells from AD donors compared to controls. Mathys and colleagues used single-nuclei sequencing and subclustered ∼2400 microglia of 48 donors into 4 subpopulations. They highlighted the Mic1 cluster as AD-pathology-associated human microglia [17]. From the 257 DAM genes investigated, only 28 were overlapping with marker genes for the Mic1 cluster and 16 of these 28 overlapping marker genes were ribosomal genes. A larger-scale investigation of microglia nuclei will be needed to reveal AD-associated microglia subpopulations in humans.

Single nucleus gene expression faithfully recapitulates cellular gene expression profiles [36,37]. Therefore, single-nucleus sequencing offers an alternative to single-cell sequencing that is especially useful in tissues from which recovering intact cells is difficult [36–38]. An important advantage of single-nucleus sequencing is the possibility to use frozen samples from brain banks containing large, well-characterized donor cohorts. For example, an improved donor cohort to differentiate the effects of natural ageing and AD pathology would be possible by comparing aged-matched (young) control donors to early-onset familial AD cases. In future, single-nucleus sequencing of microglia and including tissues of early, pre-symptomatic stages of AD will be most promising to potentially identifying (early) microglia biomarkers for AD.

## Conflict of Interest

MW, AW, SX, TM and KB were full time employees of Abbvie during the time of the studies. The remaining authors declare that the research was conducted in the absence of any commercial or financial relationships that could be construed as a potential conflict of interest.

## Author Contributions

EB, BE and SK were responsible for the overall conception of the project and provided supervision. QJ, AA, LK, conducted the experimental work, and/or analyzed the data, prepared the figures and wrote the manuscript. EG assisted with sample processing, data discussion and conducted all immunohistochemistry. NB, AM, MD, MW, AW, SX, KB, TM, YH, and TO assisted with maintenance of availability of reagents, sample processing, data discussion, optimization issues and/or data analysis. All authors contributed to the editing of the paper.

## Funding

MW, SX, AW, TM and KB are employed by AbbVie, Inc., which has subsidized the study. BE acquired funding from the foundation Alzheimer Nederland. AA, EG are funded by Abbvie. AA was supported by the Jan Kornelis de Cock-Hadders foundation. QJ was funded by the Li Ka-shing Foundation at Shantou University Medical College, China, the Abel Tasman Talent Program, University Medical Center Groningen/University of Groningen, The Netherlands, and the Graduate School of Medical Sciences (GSMS), University of Groningen, the Netherlands. LK holds a scholarship from the GSMS, University of Groningen, the Netherlands.

## Acknowledgments

We thank the Netherlands Brain Bank and the NeuroBiobank of the Institute Born-Bunge, Belgium. We thank Ortiz, T. Oshima, M. Wijering, S. Eskandar, R. van der Pijl, E. Wesseling, C. Grit and C.B. Haas for assistant and/or maintenance of sample processing and bulk and/or single cell isolations. We are grateful to W. Abdulahad, G. Mesander, T. Bijma and J. Teunis from The Central Flowcytometry Unit of the University of Groningen, University Medical Center Groningen (UMCG), for technical assistance on FACS sorting. We thank P. van der Vlies, D. Brandenburg, N. Festen and W. Uniken Venema for their assistance setting up the bc-SmartSeq2. We thank M. Meijer for ICT-related support. We appreciate sequencing-related advices provided by D. Spierings, K. Hoekstra-Wakker, J. Beenen and N. Halsema.

## Data availability

The data supporting the conclusions of this manuscript will be made available on Gene Expression Omnibus upon publication.

## Supplementary materials

**Figure S1 Donor classification based on immunohistochemistry**

(A,B) Representative immunohistochemical images of Aß and PHF-tau staining for the LPS (A) and GFS (B) of the four groups (CTR, CTR+, AD, MCI). Scale bar= 100µm. AD: clinically diagnosed Alzheimer’s Disease with Aβ plaques and/or hyperphosphorylated tau, CTR: Control, CTR+: Control with Aβ plaques and/or hyperphosphorylated tau, MCI: clinically diagnosed mild cognitive impairment with Aβ plaques and hyperphosphorylated tau.

**Figure S2 bc-Smart-seq2 single cell RNA sequencing quality of donors and clusters**

Boxplots displaying the number of detected UMIs, unique genes and mitochondrial genes per donor (A,B,D) and per cluster (D,E,F).

**Figure S3 Biological annotation of marker genes for each cluster**

Dotplots displaying the gene ratio per gene ontology term for each cluster identified in bc-Smart-seq2 analysis.

## Supplementary tables

Table S1 Detailed donor information of bulk sequenced samples.

Table S2 Detailed donor information of single-cell sequenced samples.

Table S3 Differential gene expression analysis of male vs. female bulk microglia GFS samples.

Table S4 Differential gene expression analysis of LPS vs. GFS bulk microglia samples.

Table S5 Differential gene expression analysis of enzymatic vs. mechanical dissociated bulk microglia samples.

Table S6 Cluster markers of the identified 9 clusters with bc-Smart-seq2 single cell RNA sequencing.

Table S7 Statistics of the average gene set expression changes in the clusters identified.

Table S8 Cluster markers of the identified 3 clusters of donor 2018-135 with 10X Genomics single cell RNA sequencing.

Table S9 Cluster markers of the identified 3 clusters of donor 2019-10 with 10X Genomics single cell RNA sequencing.

