## Supplementary figures and images for "Profiling microglia from AD donors and non-demented elderly in acute human post-mortem cortical tissue"

### Figure S1

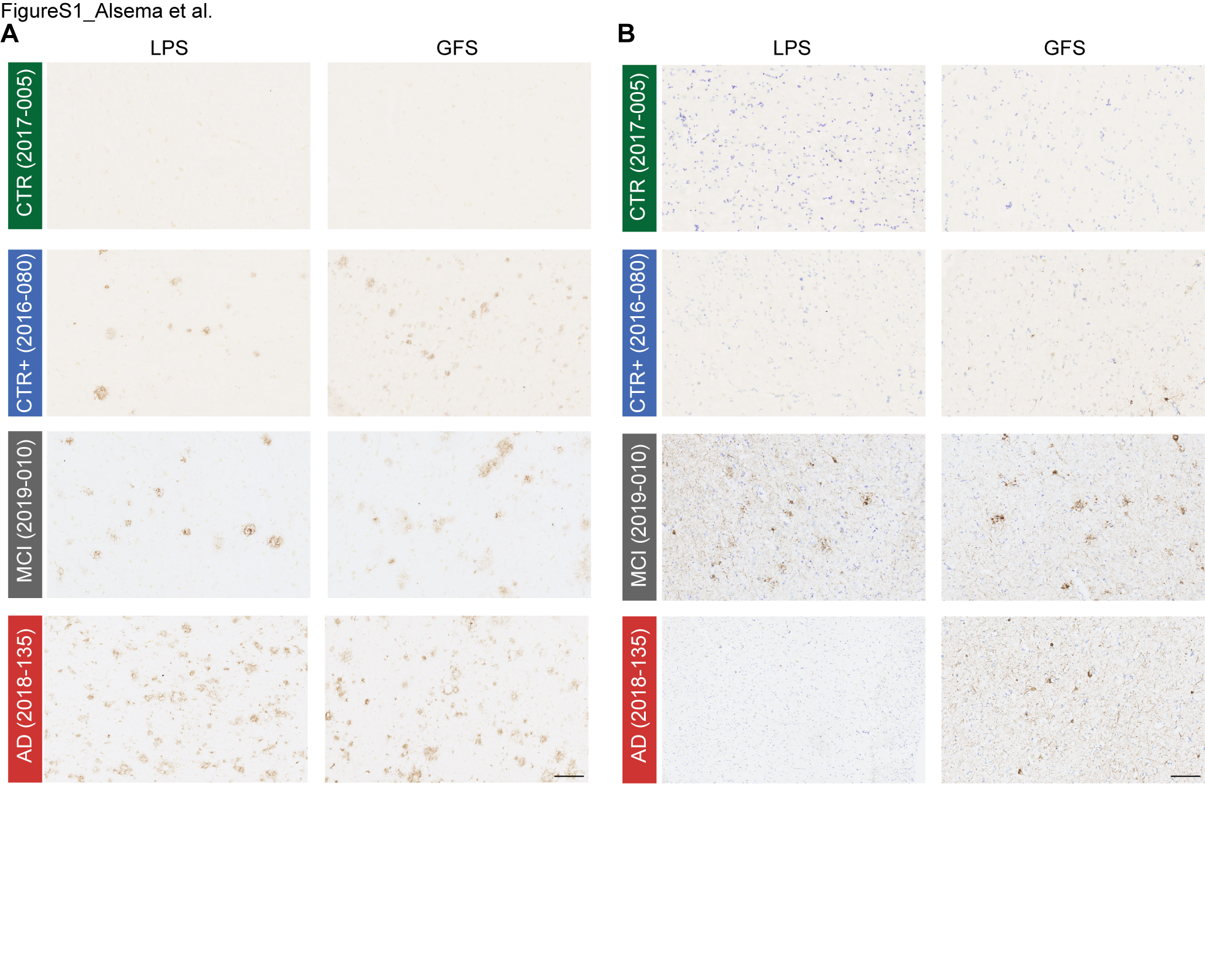

### Figure S2

FigureS2_Alsema et al.

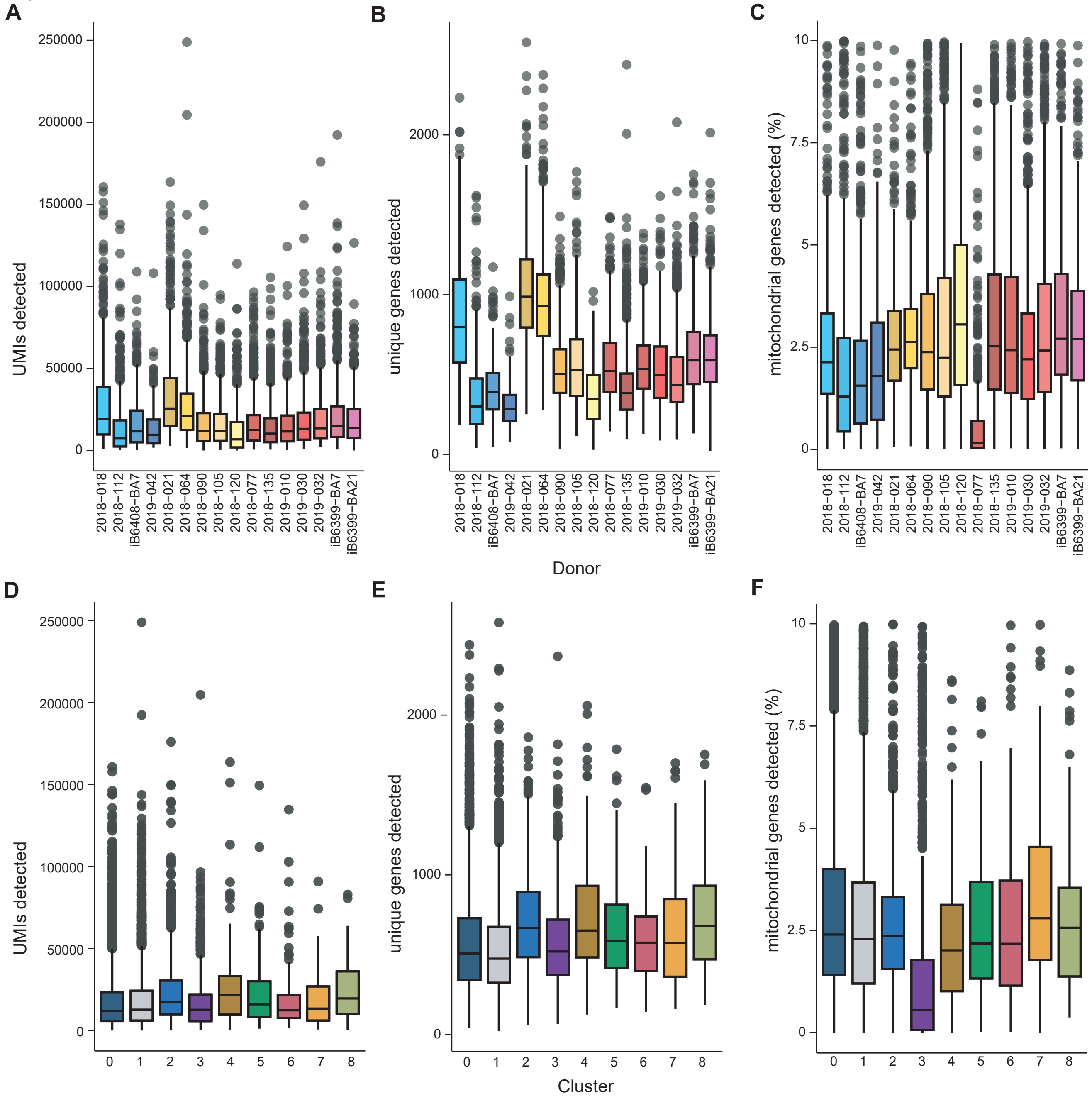

### Figure S3

FigureS3_Alsema et al.

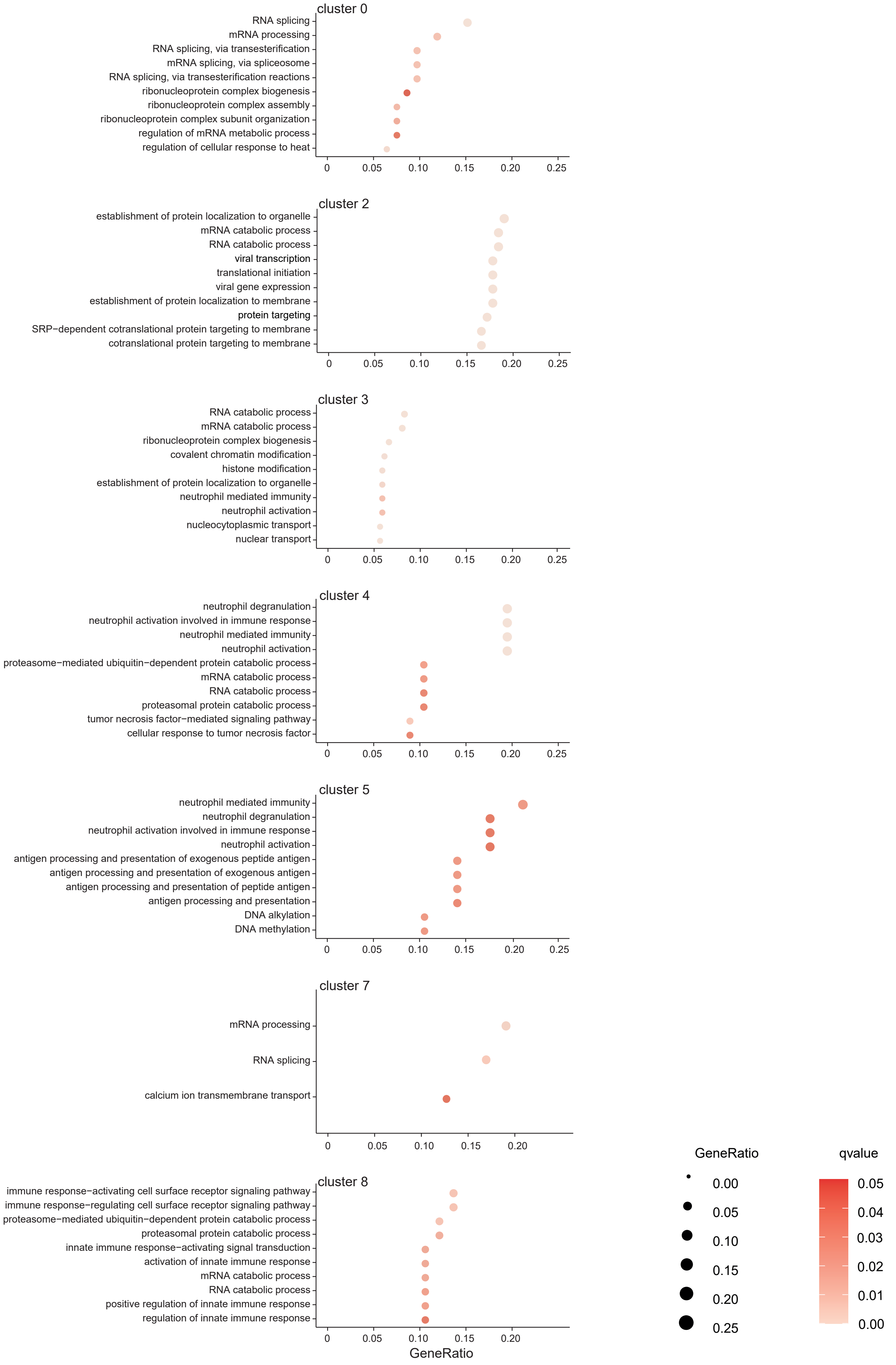
